# GeneToCN: an alignment-free method for gene copy number estimation directly from next-generation sequencing reads

**DOI:** 10.1101/2023.05.31.543012

**Authors:** Fanny-Dhelia Pajuste, Maido Remm

## Abstract

Genomes exhibit large regions with segmental copy number variation, many of which include entire genes and are multiallelic. We have developed a computational method GeneToCN that counts the frequencies of gene-specific *k*-mers in FASTQ files and uses this information to infer copy number of the gene. We validated the copy number predictions for amylase genes (AMY1, AMY2A, AMY2B) using experimental data from digital droplet PCR (ddPCR) on 39 individuals and observed a strong correlation (R=0.99) between GeneToCN predictions and experimentally determined copy numbers. We further tested the method on three different genomic regions (SMN, NPY4R, and LPA Kringle IV-2 domain). Predicted copy number distributions of these genes in a set of 500 individuals from the Estonian Biobank were in good agreement with the previously published studies. In addition, we investigated the possibility to use GeneToCN on sequencing data generated by different technologies by comparing copy number predictions from Illumina, PacBio, and Oxford Nanopore data of the same sample. Despite the differences in variability of *k*-mer frequencies, all three sequencing technologies give similar predictions with GeneToCN.

## INTRODUCTION

Copy number variation (CNV) is a type of structural variation ranging from 50 to several million base pairs (bp)^1–5^. It is an unbalanced variation where a segment of the human genome can be deleted, duplicated, or repeated multiple times and the number of repeats varies between individuals. Around 4.8 - 9.5% of the human genome contributes to CNVs^6^, a larger proportion than to SNVs, which may be associated with phenotypic traits including susceptibility to complex diseases^7–9^ especially when the copy number variation overlaps a gene region^10^. In this work, we will focus on copy number estimation of repeated genes or functional domains.

The detection of copy number differences requires a special computational approach, different from SNV analysis. The most common methods for copy number estimation from the WGS data use read depth (RD) and/or paired-end mapping (PEM)^11–16^ algorithms associated with custom-made statistical models for copy number detection. PEM-based methods map both paired ends to the reference genome and detect copy number change when the distance of mapped reads is significantly different from the mean insert size of the fragments. For this reason, PEM-based approaches cannot detect long CNVs and are not suitable for evaluating the exact copy number. Methods based on read depth use the depth of coverage information from reads mapped to the reference genome. For example, AMYCNE^17^ is an RD method dedicated specifically to studying copy numbers of the amylase gene family. RD-based methods can detect larger CNVs and can be used with paired-end reads as well as single-end reads and can also estimate more precise copy numbers. However, the accuracy may be low when estimating exact copy numbers, especially when the number of repeats is high. In addition, both approaches depend on read mapping which is time-consuming and often unreliable in complex and repetitive genomic regions. Furthermore, many methods only work using cohort data, unable to estimate copy numbers for single individuals or for a small set of samples.

An alternative approach is to use an alignment-free analysis that is based on counting and analyzing the frequencies of *k*-mers in individual genomes. *K*-mers (small substrings of DNA with length *k*) have been used for different purposes in genome analysis to efficiently handle huge amounts of genomic data^18–20^. Alignment-free methods do not require read alignment or mapping thus allowing fast and reliable genotyping of known variants^21,22^, discovering novel variants^23^ and genotyping polymorphic Alu-elements^24^. Only a handful of fully alignment-free methods have been created for estimating copy number variation of gene regions. For example, a general alignment-free CNV detection software QuicKmer2, which is also able to handle gene regions, has recently been published^25^. However, this software is paralog-specific and has difficulty handling cases where a gene or region has multiple copies in the reference genome.

In this study, we propose a novel alignment-free method GeneToCN for targeted copy number estimation of copy-variable genes. We pay special attention to the selection of robust and reliable *k*-mers in gene regions. Our approach allows estimating copy numbers for individual samples without the requirement of cohort data. We demonstrate our method’s accuracy on the amylase gene family as well as general useability on three other genes (NPY4R, SMN, and LPA Kringle IV type 2 domain).

## RESULTS

### Method for alignment-free gene copy number estimation

The working principle of the GeneToCN method is the following. First, a custom database is created consisting of carefully selected *k*-mers a) from a gene region and b) from the flanking regions of the same gene. The flanking regions are used to estimate the local depth of coverage (DOC), which is used as a reference in copy number estimation. The choice of representative *k*-mers for each gene is a crucial step of our method. To select the most robust and reliable set of reference *k*-mers, we apply several filters based on their uniqueness in the reference genome and their GC-content (described in Methods). The *k*-mer selection process is automated with GeneToKmer scripts (Figure 1).

**Figure 1.**
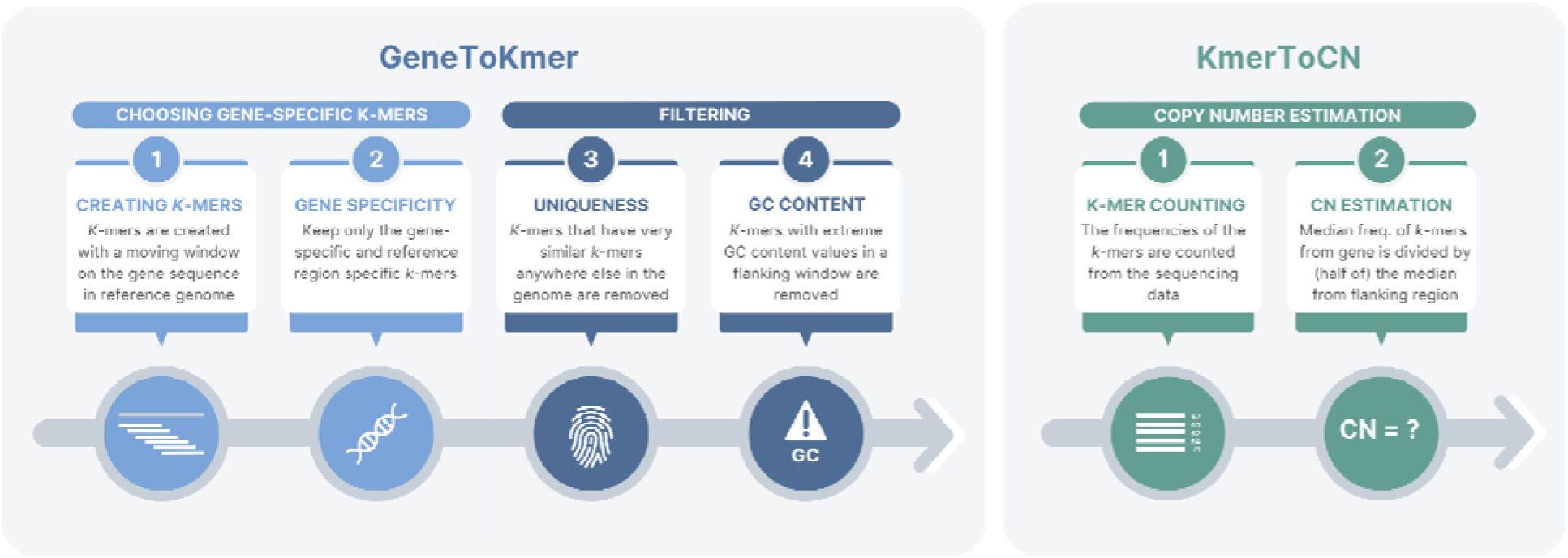
Overview of the method

Copy number estimation in each studied individual starts with counting the frequencies of the selected gene-specific *k*-mers directly from the raw sequencing reads of this individual. The copy number of each gene is calculated by dividing the median frequency of gene-region *k*-mers by the median frequency of flanking-region *k*-mers and multiplying by 2 (the ploidy of the human genome). The resulting copy number is decimal, but it can be rounded to the nearest integer if an integer copy number is preferred/required for interpretation. In this article, we use decimal numbers for correlation analysis and integers for concordance analysis.

Our method has a unique approach for handling regions in the reference genome that have multiple copies. For example, AMY1 is present in 3 copies in the reference. Unlike other methods that generally estimate the copy number separately for each of these copies, our GeneToKmer script has the flexibility to either treat them separately or to define all 3 copies as a single gene. In the first case, we use the *k*-mers specific to each different copy, whereas in the latter case, we use only *k*-mers that are present in all 3 copies. By avoiding the use of *k*-mers that may be variable due to recent mutations and are not present in all copies of a given gene, we can improve the accuracy of copy number predictions.

### Copy number estimation in AMY1, AMY2A, and AMY2B gene regions

First, we investigated the performance of the GeneToCN method using the well-studied alpha-amylase gene family^26^. Amylase is a digestive enzyme that catalyzes the hydrolysis of starch and is present in human saliva as well as in the pancreas. The human reference genome has three copies of the salivary amylase gene AMY1 and one copy of the pancreatic amylase genes AMY2A and AMY2B. There is also a pseudogene AMYP1 containing a large part of the sequence of AMY2A. The copy numbers of amylase genes are highly variable, especially for AMY1, for which it varies from 2 to 22^27–29^. The copy number of AMY2A varies from 0 to 8, the least copy-variable is AMY2B with a copy number from 2 to 6. For testing the GeneToCN method, the *k*-mers were selected for each of these amylase genes and copy numbers were estimated from Illumina sequencing reads of 500 individuals from the Estonian Biobank (EstBB). Although the frequency of individual *k*-mers is variable, the consolidated information from all gene-specific *k*-mers allows reliable detection of differences between flanking region and gene region (Figure 2).

**Figure 2.**
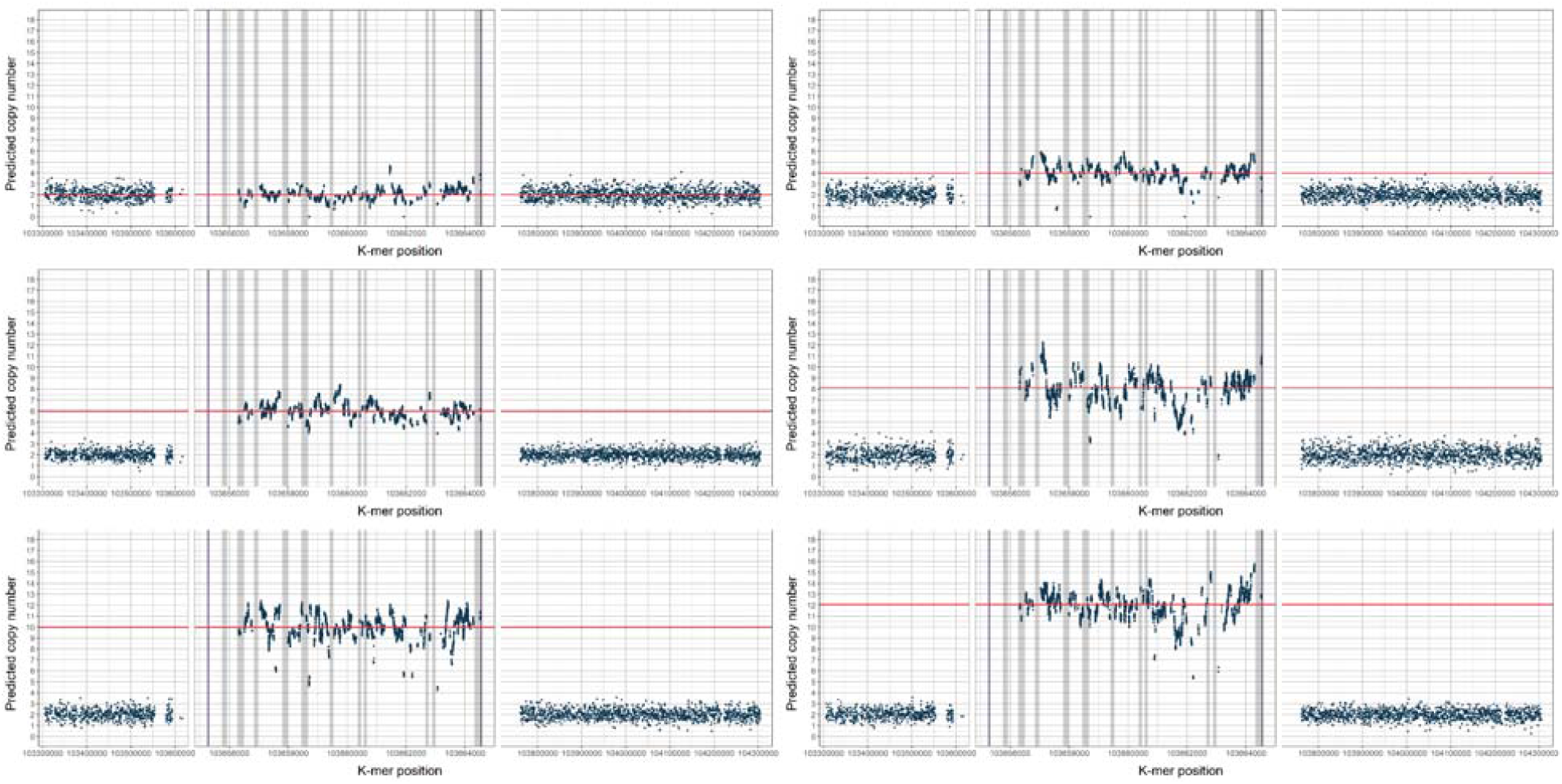
Normalized *k*-mer frequencies (y-axis) in different individuals, predicted to have 2, 4, 6, 8, 10, and 12 copies of the AMY1 gene. The x-axis shows the *k*-mer locations on chromosome 1. The horizontal red line marks the copy number estimated by GeneToCN. Each panel shows a 5’-flanking region, a zoomed-in AMY1 gene region, and a 3’-flanking region. Exon regions are shown in grey.

The distributions of predicted copy numbers in 500 individuals are shown in Figure 3. As shown in previous studies^29,30^, even copy numbers were more common than odd copy numbers for the AMY1 gene, with 73% of studied individuals having 4, 6, 8, or 10 copies. AMY2A copy numbers varied from 0 to 5 and AMY2B copy numbers varied from 2 to 4, which is also consistent with distributions observed in previous studies^29,30^. For AMY2B we observed a duplication breakpoint within the first third of the gene at position Chr1:103,561,000 (Suppl. Figure S1). For those individuals copy number 4 was called by GeneToCN.

**Figure 3.**
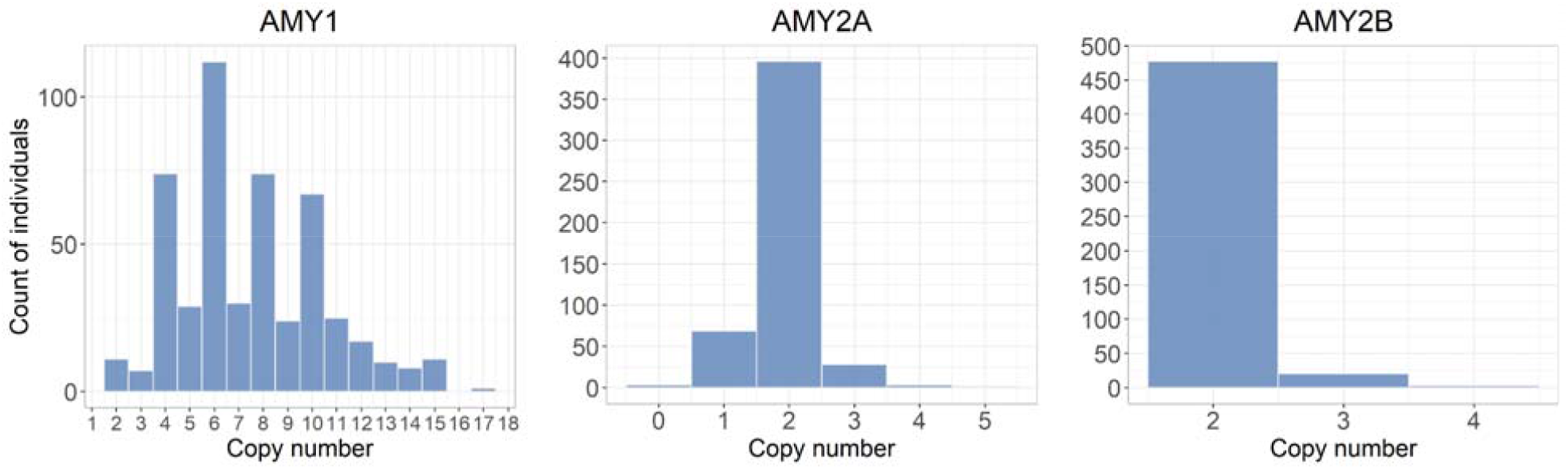
Amylase copy number distributions in 500 Estonian individuals from EstBB.

Previous studies on amylase gene copy numbers have shown that 98% of individuals have the same parity of the copy numbers of AMY1 and AMY2A, meaning that the copy numbers of these genes are usually both either even (more frequent) or both odd (rarely) at the same time^29^. GeneToCN predictions from 500 EstBB individuals (Figure 3) showed the same parity in 85% of tested individuals, which is significantly higher parity than expected by chance alone (P=1.253e^-14^), assuming that the AMY1 and AMY2A alleles are randomly paired.

### Experimental validation

In addition to the analyses of frequency distributions (Figure 3), the GeneToCN method was also validated experimentally using data from digital droplet PCR (ddPCR). For this, we used 40 individuals from EstBB, for which we had copy number data of AMY genes from previously published ddPCR experiments^29^. Although the experimental methods do not guarantee 100% accurate results, ddPCR has been used as the gold standard for experimental copy number determination^31,32^ and is thus a good reference for the evaluation of the GeneToCN method. The correlation of copy number estimates from GeneToCN and ddPCR experiments for these 40 individuals are shown in Figure 4A. Only one individual had a difference larger than 1 copy (8 copies by ddPCR and 12 copies predicted by GeneToCN). We examined the *k*-mer frequency plot of this individual (Suppl. Figure S2), but could not detect any reasons that could explain the difference in predictions in this individual. All *k*-mers in the gene region support the prediction of 12 copies without any regional fluctuation. This data point was excluded from further calculations, as a likely human error.

**Figure 4.**
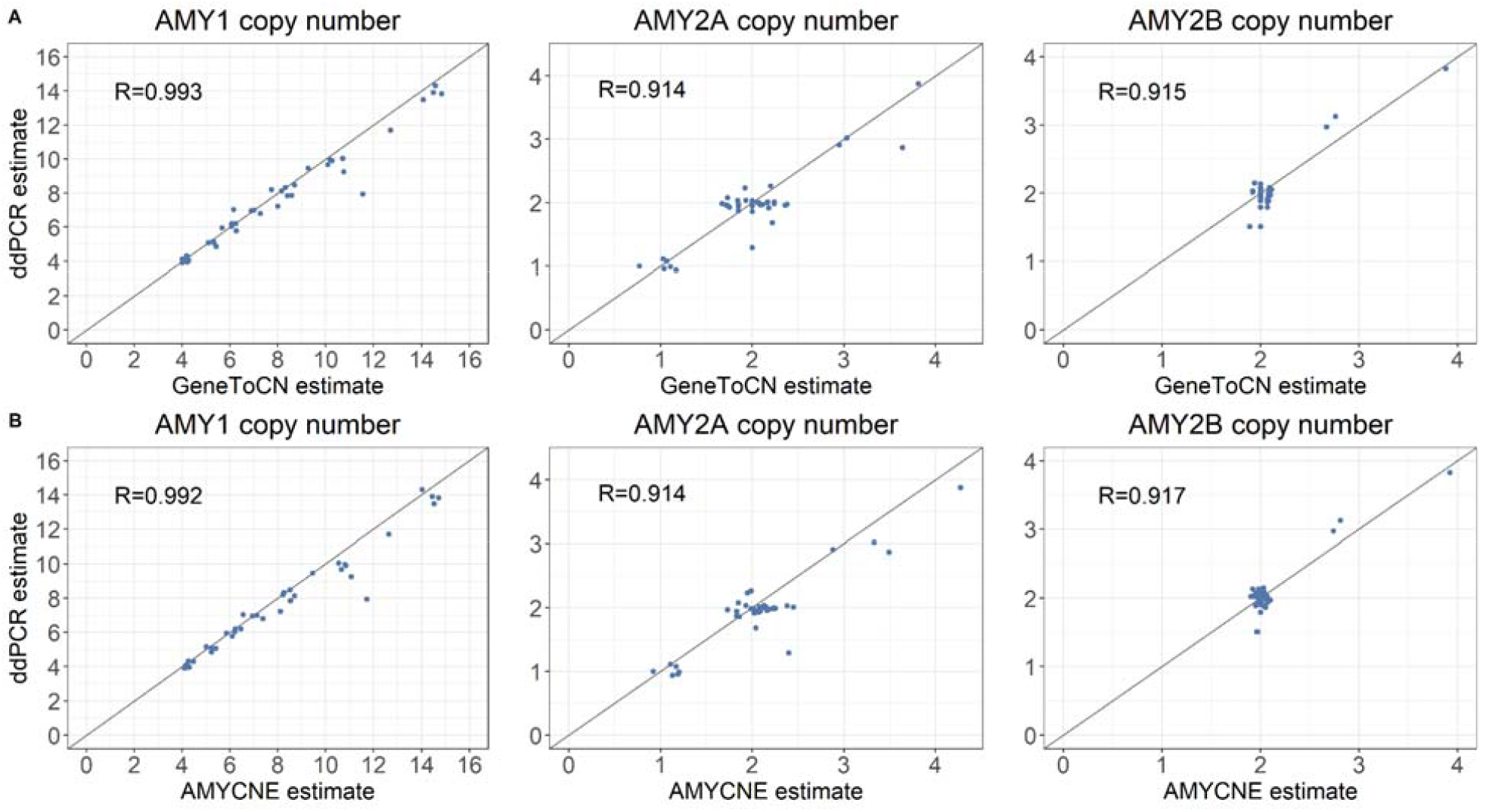
Correlation between copy number estimates from GeneToCN and ddPCR (A) and from AMYCNE and ddPCR (B) using 40 EstBB samples.

For numerical comparison, the correlation coefficient R was calculated from the raw results of both methods and shown as a decimal number, whereas the concordance was calculated based on the integer copy number values (raw result rounded to the nearest integer). The results for all three amylase genes are shown in Supplementary Table S1. The correlation coefficient between predictions and experimental results was 0.91 for AMY2A and AMY2B and 0.99 for AMY1. The concordance was 74%, 97% and 100% for AMY1, AMY2A and AMY2B, respectively.

We observed that the correlation coefficient between GeneToCN and ddPCR predictions is 0.99, but the concordance of integer predictions is only 74%. This is caused by the tendency of GeneToCN to slightly overestimate the copy number in individuals with >8 copies of AMY1 (Figure 4). Predictions could be improved, for example by using linear regression. We were able to increase the concordance of GeneToCN predictions to 85% by using the regression formula y = 0.9622*x for the correction. However, we did not implement this correction in the GeneToCN code because we do not have an independent dataset for testing the robustness of the correction on other gene regions.

### Comparison with AMYCNE

For comparison, the copy numbers of AMY genes in the same individuals were also estimated using a previously published software AMYCNE^17^. AMYCNE uses an algorithm based on read mapping and subsequent read depth analysis for copy number estimation. AMYCNE has been previously validated on amylase genes and would therefore be expected to be optimized for the analysis of these genes. In correlation analysis with ddPCR, we observed comparable accuracy for both KmerToCN and AMYCNE (Figure 4B). For integer copy numbers, the predictions of AMY2A and AMY2B gene copy numbers were comparable, whereas GeneToCN predictions for the AMY1 gene had higher concordance with ddPCR results (Table S1). The parity of AMY1 and AMY2A copy number predictions in 39 individuals were 87%, 82% and 67% for ddPCR, GeneToCN and AMYCNE, respectively.

### Testing on different gene regions

We tested GeneToCN thoroughly on the amylase gene region. However, it would be important to know if the same method can be used for the estimation of copy numbers of other genes, particularly whether a sufficient number of *k*-mers can be selected from gene regions and flanking regions. We created custom *k*-mer databases for a set of genes from different genomic regions and with different copy numbers: survival of motor neuron genes SMN1 and SMN2^33–35^, the human pancreatic polypeptide receptor gene NPY4R^36,37^, and the LPA gene. In the latter case, the repeated region consists of only one protein domain, the 5.5 kb long Kringle-IV type 2 domain, which spans over 2 exons^38,39^. This case allowed us to validate the suitability of the method not only on full genes but on shorter high-copy repeats as well. For these genes, we tested whether an adequate number of *k*-mers can be selected and whether their distributions of predicted copy numbers coincide with previously published copy number distributions.

For each of these genes, *k*-mers were selected with GeneToKmer, and copy numbers were estimated with the KmerToCN tool from the 500 EstBB individuals as described above. The distributions of predicted copy numbers are shown in Figure 5. The copy numbers for the NPY4R gene varied from 2 to 8 and the most common copy number was 4. The copy numbers for SMN1 varied between 1 and 3, whereas for the SMN2 gene, the copy number estimates were between 0 and 3. For the Kringle-IV type 2 domain, the copy numbers were between 18 and 58 (mean 39.2), with the most common copy number being 40. In a previous study, where copy numbers of the Kringle-IV type 2 region were estimated with the Genome STRiP for a larger sample of 2284 Estonians from the EstBB, the copy numbers varied between 12 and 63 with a mean of 39.7^40^. These results, particularly the fact that mean copy numbers of the Kringle-IV type 2 domain are very similar, confirm that the GeneToCN method is robust and usable for the estimation of copy numbers for even high-copy repeats.

**Figure 5.**
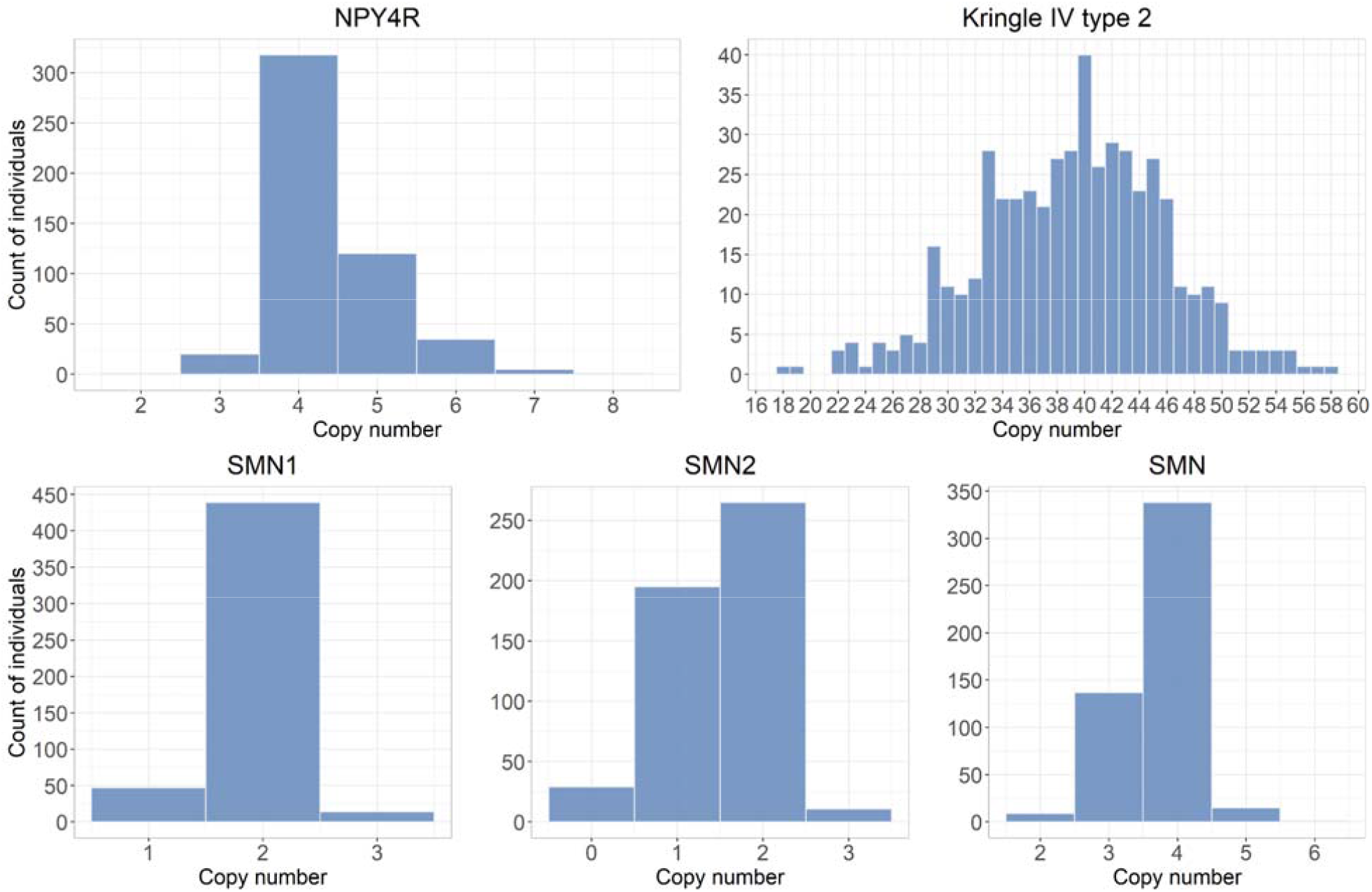
Copy number distributions for 500 individuals, estimated using KmerToCN. Panel SMN represents the sum of SMN1 and SMN2 gene copies.

Analysis of SMN1 and SMN2 genes revealed some limitations of the GeneToCN method. These two genes have a nucleotide-level identity of about 99.9%, therefore only a limited number of gene-specific *k*-mers (268 and 267, respectively) could be selected. It is not clear how accurately copy numbers are inferred from such a small number of *k*-mers. To better evaluate the accuracy of SMN1 and SMN2 copy numbers, a separate *k*-mer database was created for estimating the total number of SMN copies, using the *k*-mers present in both genes. For 77% of the individuals, the sum of SMN1 and SMN2 copy numbers estimated separately matched the total SMN copy number. The 500 individuals were then divided into two groups based on whether the sum matched the total SMN copy number or not. We observed that the group where the copy numbers did not match had significantly lower (Wilcoxon test, P=2.2*10^−16^) copy number values for SMN2, as well as for SMN1 (P=0.0023). This can be explained by single nucleotide variants in the SMN genes that may cause underestimation of the SMN2 and in some cases SMN1 copy number. Overall, it seems that the number of gene-specific *k*-mers in SMN1 and SMN2 is too small to allow reliable estimation of their copy numbers separately. However, both SMN genes together had >16,000 gene-specific *k*-mers allowing reliable prediction of their cumulative copy number.

### Copy numbers estimated from long-read sequencing data

Long-read sequencing data from Oxford Nanopore and PacBio sequencing technologies were used in addition to Illumina reads to evaluate how the method works on other sequencing data apart from Illumina. The comparisons were done on a reference sample CHM13 that has been sequenced by three different technologies^41^ to 50x (Illumina), 120x (Oxford Nanopore), and 30x (PacBio) depth of coverage. The copy numbers were estimated for all six gene regions and the results are shown in Supplementar Table S2. Overall, the copy number predictions are similar with all three technologies, fluctuating within 10% from each other.

A visual overview of *k*-mer frequency variation from AMY1, AMY2A, and AMY2B regions is shown in Figure 6, and NPY4R, SMN, and LPA Kringle IV-2 regions are shown in Supplementary Figure S3. We observed a difference in variations of *k*-mer frequencies between the three technologies. The Oxford Nanopore data is affected by the high mutation rate, resulting in high variability. PacBio data is the least variable. However, these differences do not have any systematic adverse effects on the copy number estimation, making us conclude that all three technologies are suitable for the alignment-free inference of copy numbers.

**Figure 6.**
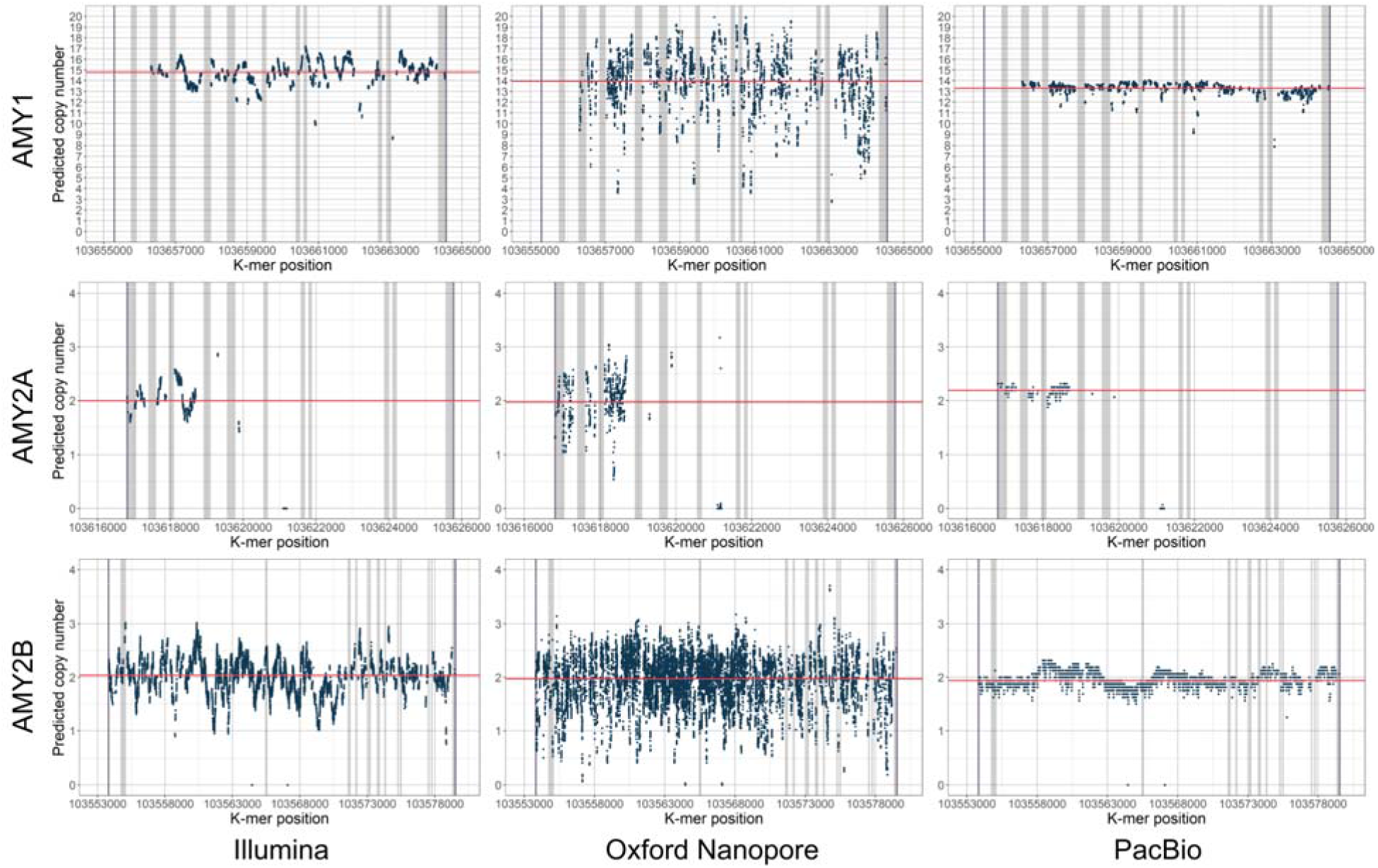
*K*-mer frequencies in AMY1, AMY2A, and AMY2B regions from sequencing data generated by Illumina, Oxford Nanopore, and PacBio technologies.

## METHODS

### Creation of *k*-mer databases with GeneToKmer

The *k*-mer databases for gene regions were compiled using the GeneToKmer program that utilizes tools from version 4.2.16 of the GenomeTester4 package from GitHub^20^. The *k*-mer length used throughout this study was 25. The coordinates used for each region can be seen in Supplementary Table S3. The first step of the *k*-mer selection process is creating a set of overlapping *k*-mers for each gene region. These *k*-mers were generated by a moving window using the GListMaker and human reference genome GRCh38p10. For regions that had multiple copies in the reference genome (for example Kringle IV type 2 with 6 copies or AMY1 with 3 copies), separate *k*-mer lists were initially created for each region and then the intersect of the common *k*-mers was taken.

For the selection of region-specific and unique *k*-mers, the frequencies of all the *k*-mers were then counted from the reference genome. For this, we used a total *k*-mer list compiled from the GRCh38p10 reference genome. The frequencies of all *k*-mers in the genome were obtained using the tool GListQuery from GenomeTester4. *K*-mers were considered unique and gene-specific when the frequency in the reference genome was equal to the number of copies of that gene in the reference (for example 3 for AMY1). We used the 1 mismatch (-mm 1) option to exclude any *k*-mers that had identical *k*-mers or *k*-mers with edit distance 1 present somewhere else in the reference genome (except for SMN1 and SMN2 regions where we did not use that option). The region-specific *k*-mers were further filtered by GC-content. We filtered out all *k*-mers with a GC-content value lower than 20 or higher than 65.

It is possible to use GeneToKmer in the same manner for compiling databases for flanking *k*-mers. However, in regions with repeated content, the flanking region might need to be rather large (several million base pairs) and finding unique *k*-mers from these regions might require lengthy calculations. Alternatively, a reference region with a non-variable copy number may be chosen further away from the gene. In this project, we decided to use subsets of unique *k*-mers previously compiled for NIPTmer prenatal diagnostic software^42^. As CHM13 assembly was used in this project, we further filtered the NIPTmer *k*-mers based on the uniqueness of these *k*-mers in the CHM13 assembly. The coordinates of used regions and the number of selected gene-specific and flanking *k*-mers are shown in Supplementary Table S3.

### Copy number estimation

Copy number estimations were done with KmerToCN software scripts. These use either gmer_counter from the FastGT toolkit^22^ or optionally GListMaker and GListQuery from the GenomeTester4 toolkit^20^ for *k*-mer counting, depending on the input file type of the sequencing data (FASTQ or *k*-mer list).

AMYCNE required running another tool called TIDDIT, both tools are written in Python. AMYCNE needed some modifications in the code as well as in several input files to work. For AMY2A, the sequence coordinates proposed by the authors were altered by keeping only the first part of the gene up until the fourth exon (since the rest of the gene sequence is identical to the sequence of the pseudogene AMYP1), which improved the overall correlation with ddPCR results from 0.54 to 0.92 and integer copy number concordance from 0.58 to 0.98.

The ddPCR copy numbers used for validation of the method and the process of ddPCR experiments were described in a previous study^29^.

### Sequencing data

Method validation and copy number estimations were conducted using 500 samples from the Estonian Biobank, of which 40 samples were used also for the comparison of AMY1, AMY2A, and AMY2B copy numbers estimated by different methods. The Illumina sequencing data (ca 30x depth of coverage, read length of 151 bp) for the Estonian Biobank samples were retrieved from the Estonian Genome Centre. The Illumina, Oxford Nanopore, and PacBio sequencing data for CHM13^41^ were retrieved from https://github.com/marbl/CHM13/blob/master/Sequencing_data.md.

### Computational performance

Creating the *k*-mer databases for a gene region with GeneToKmer typically takes less than 15 minutes. The time usage depends mostly on the length of the gene regions. KmerToCN can process one individual in 30-60 minutes if the source file is a FASTQ file. This is similar to the speed we have demonstrated previously for other alignment-free genome analysis tools^22,24^. The minimum amount of required RAM depends on the overall number of *k*-mers used to describe repeated gene regions. Although we did not perform an analysis of memory usage in this particular work, our previous studies suggest that the frequencies of 30 million SNV-specific *k*-mers can be counted with as little memory as 8GB^22^. The performance was measured on a Linux server with 64 CPUs (2.27 GHz) and 512GB RAM.

## DISCUSSION

In this study, we propose a novel alignment-free method GeneToCN for targeted gene copy number estimation. In this approach, we use local *k*-mer frequencies from the flanking regions of a gene as a reference for normalization. Defining the “flanking regions” for *k*-mer selection assumes that we know the approximate breakpoints of the copy-variable region, which in most cases are already available from previous studies. Alternatively, *k*-mers from other known non-copy-variable regions, preferably located near the targeted genes, can be used. Novel breakpoints can be detected from *k*-mer frequency plots of each individual.

What are the advantages of using raw sequencing reads instead of mapped reads for variant detection, particularly for gene copy number estimation? The sequencing data are often stored in a BAM or CRAM format where reads are already mapped to the reference genome. However, there are some important benefits for variant detection and copy number estimation directly from raw sequencing reads. Most importantly, using the raw data makes the method more robust and easy to use. The mapping software requires experienced users to choose the right parameter values. An alignment-free method averts the effect of methodological errors in read mapping (due to mismapping or incorrect reference sequence) which simplifies the analysis process and may increase the accuracy of the results in some regions. Also, speed and the consequent decrease in computational costs are beneficial in large-scale studies where thousands of individuals need to be analyzed. For example, in a meta-analysis of large datasets, it is necessary to use the same analysis pipeline for all individuals. In this case, it might not be practical to re-map all the reads in meta-analysis projects, but rather use a fast alignment-free approach.

What is the minimum depth of coverage for alignment-free analysis? This is a complicated question without an easy answer. There is an interplay between the depth of coverage and number of k-mers in the region. It is assumed that a higher overall number (either because of sequencing depth or from region width) of k-mers would give more accurate predictions. Another factor is how equally the k-mers are distributed over the gene region. A closely located (or overlapping) set of k-mers is more prone to be influenced by local fluctuations in frequency and therefore less reliable. In a previous study, we conducted an in-depth analysis suggesting that 20 is the minimum required depth of coverage for alignment-free genotyping of single nucleotide variants^22^. For copy number predictions we use hundreds or thousands of *k*-mers (for example 3095 for AMY1), therefore a smaller depth of coverage might be sufficient. However, the exact limits of the method and correlation between the number of k-mers per region, depth of coverage and accuracy of copy number predictions need further investigation.

How to explain the differences between the two computational methods GeneToCN and AMYCNE^17^? In most analyses, they demonstrate very similar performance. For example, their correlation with experimental ddPCR predictions was nearly identical (Table S1). The difference appeared only in copy number prediction of the AMY1 gene, which has up to 16 copies. For the AMY1 region, GeneToCN had higher concordance with ddPCR results (74% vs 67%) and a higher level of parity between AMY1 and AMY2A copy numbers (82% vs 67%) compared to AMYCNE. This difference in high copy number predictions could appear from the different approaches we use for filtering *k*-mers in regions where a gene is represented in multiple copies in the reference genome.

As a further development of GeneToCN, we plan to compile and publish *k*-mer databases for all genes that are copy-variable or contain smaller copy-variable regions of interest. This would provide users with an easily accessible toolbox for alignment-free copy-number prediction.

## DATA AVAILABILITY

The source code and *k*-mer databases for analyzed genes are available on GitHub (https://github.com/bioinfo-ut/GeneToCN). The binaries and source code of the *k*-mer counting software GenomeTester4 are available on GitHub (https://github.com/bioinfo-ut/GenomeTester4/). GenomeTester4 is distributed under the terms of GNU GPL v3, and the *k*-mer databases are distributed under the Creative Commons CC BY-NC-SA license.

## ACKNOWLEDGMENTS

This work was funded by institutional grant IUT34-11 from the Estonian Ministry of Education and Research, grant SP1GVARENG from the University of Tartu, and the EU ERDF grant No. 2014-2020.4.01.15-0012 (Estonian Centre of Excellence in Genomics and Translational Medicine). The cost of the NGS sequencing of the individuals from the Estonian Genome Centre was partly covered by the Broad Institute (MA, USA) and the PerMed I project from the TERVE program. The genome data was collected and used with ethical approval Nr. 206T4 (obtained for the project SP1GVARENG). The computational costs were partly covered by the High-Performance Computing Centre at the University of Tartu.

The authors thank Steven A McCarroll and Reedik Mägi for sharing the copy number data from ddPCR experiments, Tarmo Puurand for creating the 25-mer lists for EstBB individuals, and Lauris Kaplinski for the help with the NIPTmer *k*-mer set. We thank Tarmo Puurand and Maarja Jõeloo for the critical reading of the manuscript.

## AUTHOR CONTRIBUTIONS

F.D.P. collected all the data, wrote the software, performed all analyses, and participated in writing the manuscript. M.R. conceived the idea, supervised the work, and participated in writing the manuscript.

