## Supplementary material for "GeneToCN: an alignment-free method for gene copy number estimation directly from next-generation sequencing reads"

**SUPPLEMENTARY DATA**

**Table S1.** Comparison of the results from GeneToCN and from AMYCNE to the experimental results from digital droplet PCR

| | *Correlation coefficient R* | | *Concordance* | |
|---|---|---|---|---|
| | *KmerToCN* | AMYCNE | *KmerToCN* | AMYCNE |
| *AMY1* | 0.993 | 0.992 | 74.4% | 66.7% |
| *AMY2A* | 0.914 | 0.914 | 97.4% | 97.4% |
| *AMY2B* | 0.915 | 0.917 | 100% | 100% |

* This result was achieved after manually removing the part that is repeated in pseudogene AMYP1. Without the removal of the problematic region, the AMYCNE gave a correlation coefficient of R=0.54 and a concordance of 58%.

**Table S2.** Copy numbers of different genes estimated for the CHM13 cell line from Illumina, Oxford Nanopore, and PacBio sequencing data.

| *Gene region* | *Illumina* | *Nanopore* | *PacBio* |
|---:|---:|---:|---:|
| *AMY1* | 14.8 | 14.0 | 13.3 |
| *AMY2A* | 2.0 | 2.0 | 2.2 |
| *AMY2B* | 2.0 | 2.0 | 2.0 |
| *NPY4R* | 3.8 | 4.6 | 4.5 |
| *SMN* | 3.9 | 4.4 | 3.7 |
| *LPA Kringle IV-2* | 45.2 | 39.9 | 48.7 |

**Table S3.** The coordinates of gene-specific regions for *k*-mer selection, used in this study. The reference genome was GRCh38p10.

| Region | Chr | Region coordinates | Number of gene region *k*-mers | Flanking region coordinates | Number of flanking region *k*-mers |
|---|---|---|---|---|---|
| AMY1 | 1 | 103,655,290 –<br>103,664,554<br>103,687,415 –<br>103,696,680<br>103,750,406 –<br>103,758,690 | 3095 | 103,305,000 -<br>104,305,000 | 1875 |
| AMY2A | 1 | 103,616,811 –<br>103,625,780 | 738 | | |
| AMY2B | 1 | 103,553,815 –<br>103,579,534 | 14764 | | |
| NPY4R | 10 | 46,461,099 –<br>46,465,958<br>47,918,662 –<br>47,923,524 | 4042 | 46,781,000 -<br>48,474,000 | 2099 |
| LPA Kringle IV-2 | 6 | 160,639,460 –<br>160,645,006<br>160,633,913 –<br>160,639,459<br>160,628,367 –<br>160,633,912<br>160,622,823 –<br>160,628,366<br>160,617,277 –<br>160,622,822<br>160,611,722 –<br>160,617,276 | 1781 | 160,647,000 -<br>161,617,000 | 2456 |
| SMN1 | 5 | 70,925,030 -<br>70,953,942 | 268 | 69,210,000 -<br>69,534,000<br>71,324,000 -<br>72,211,000 | 2633 |
| SMN2 | 5 | 70,049,638 -<br>70,078,522 | 267 | | |
| SMN | 5 | 70,925,030 -<br>70,953,942<br>70,049,638 -<br>70,078,522 | 16273 | | |

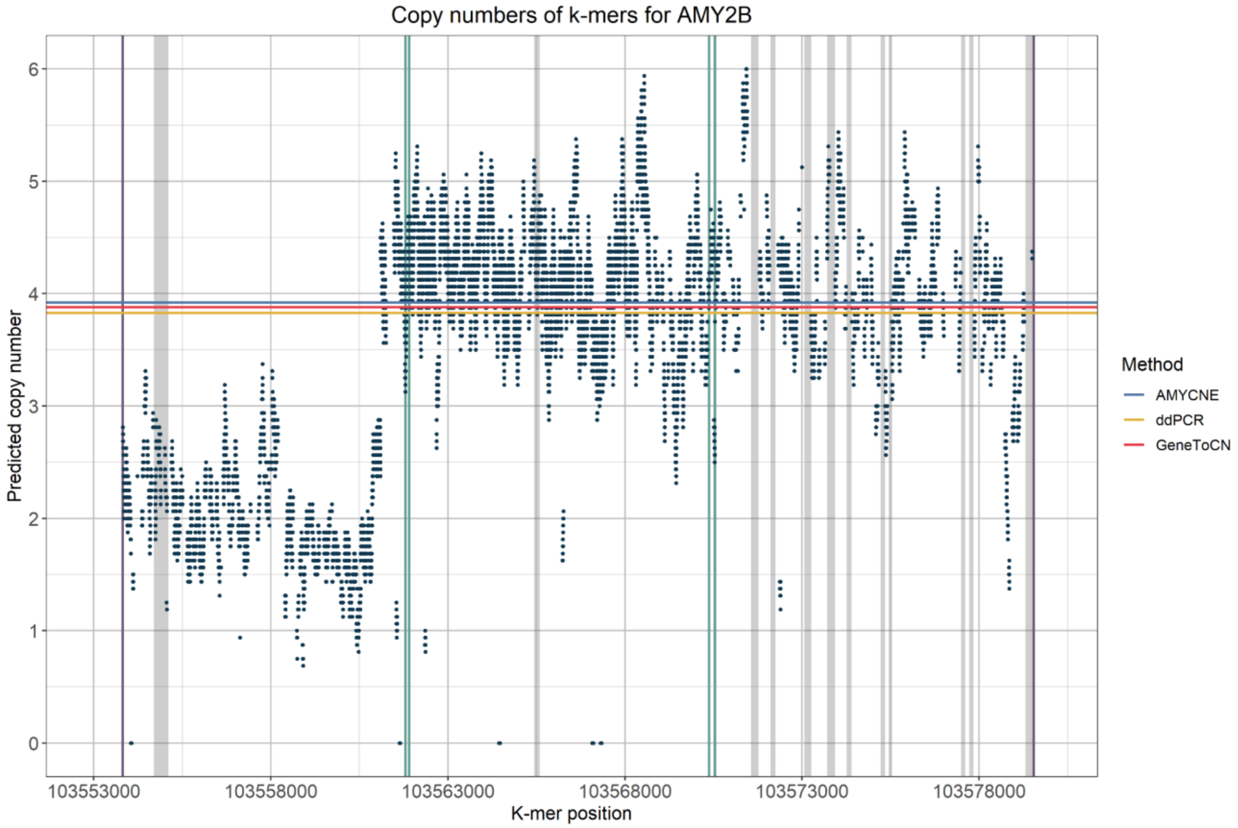


**Figure S1.** An example of atypical copy number change within the gene (atypical breakpoint). The x-axis shows the *k*-mer location in chromosome 1. The horizontal red line marks the average copy number estimated by KmerToCN. Green lines denote the locations of ddPCR primers.

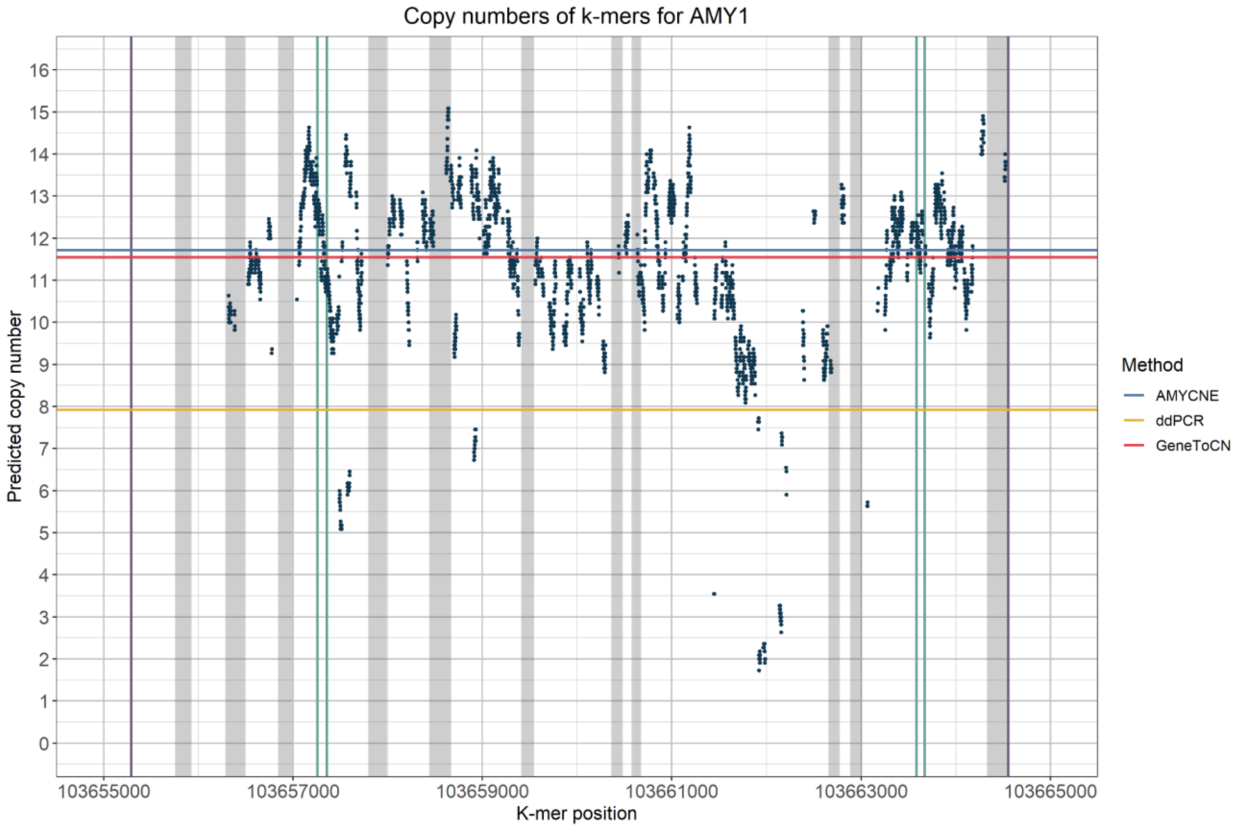


**Figure S2.** An individual with a large difference between ddPCR and GeneToCN predictions. Green vertical lines indicate the locations of ddPCR primer pairs. We can observe that k-mer frequencies in PCR primer regions that influence the ddPCR predictions have no obvious differences from the copy number estimated by GeneToCN and AMYCNE.

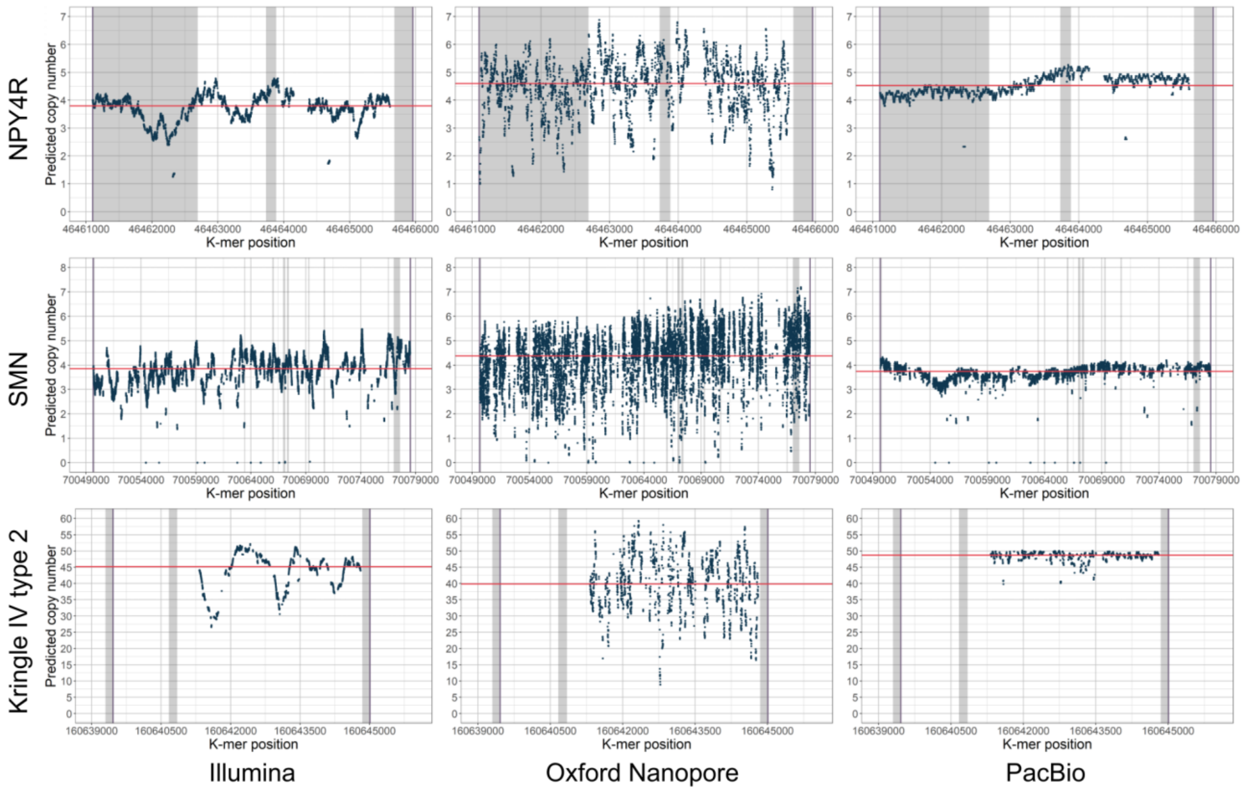


**Figure S3**. *K*-mer frequencies in NPY4R, SMN, and LPA Kringle IV regions from sequencing data generated by Illumina, Oxford Nanopore, and PacBio technologies.
